# Biased activation of the receptor tyrosine kinase HER2

**DOI:** 10.1101/2022.12.04.519064

**Authors:** Claudia Catapano, Johanna V Rahm, Marjan Omer, Laura Teodori, Jørgen Kjems, Marina S Dietz, Mike Heilemann

## Abstract

HER2 belongs to the ErbB sub-family of receptor tyrosine kinases and regulates cellular proliferation and growth. Different from other ErbB receptors, HER2 has no known ligand. Activation occurs through heterodimerization with other ErbB receptors and their cognate ligands. This suggests several possible activation paths of HER2 with ligand-specific, differential response, which so far remained unexplored. Using single-molecule tracking and the diffusion profile of HER2 as a proxy for activity, we measured the activation strength and temporal profile in live cells. We found that HER2 is strongly activated by EGFR-targeting ligands EGF and TGFα, yet with a distinguishable temporal fingerprint. The HER4-targeting ligands EREG and NRGβ1 showed weaker activation of HER2, a preference for EREG, and a delayed response to NRGβ1. Our results indicate a selective ligand response of HER2 that may serve as a regulatory element. Our experimental approach is easily transferable to other membrane receptors targeted by multiple ligands.

**Highlights:**

- HER2 exhibits heterogeneous motion in the plasma membrane
- The fraction of immobile HER2 correlates with phosphorylation levels
- Diffusion properties serve as proxies for HER2 activation
- HER2 exhibits ligand-specific activation strength and temporal profiles

**Graphical Abstract:** 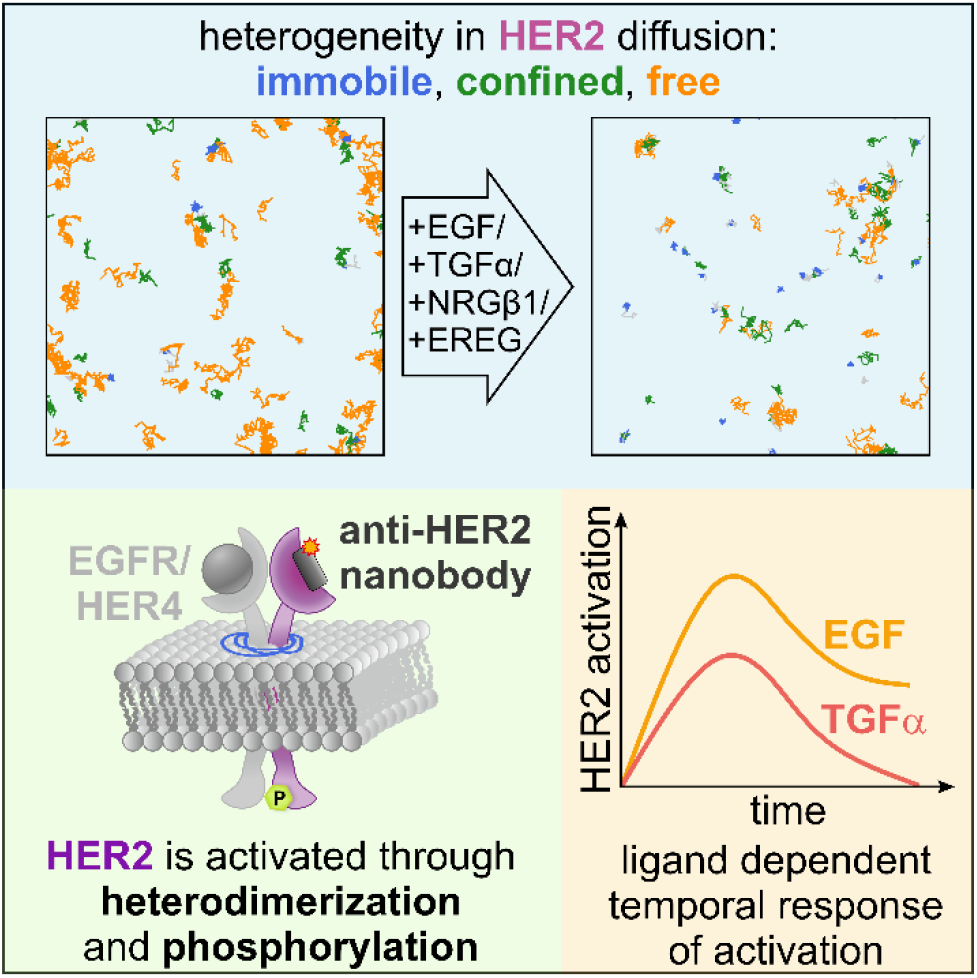

## Introduction

Receptor tyrosine kinases (RTKs) play a pivotal role in a multitude of fundamental processes such as proliferation, differentiation, and migration (Casaletto and McClatchey 2012). They are classified into numerous subfamilies based on functional or structural similarities, where each receptor group binds characteristic growth factors (Trenker and Jura 2020; Ullrich and Schlessinger 1990). Growth factor binding to receptors initiates cellular signal transduction by inducing RTK dimerization, tyrosine phosphorylation, and recruitment of downstream signaling proteins (Casaletto and McClatchey 2012; Du and Lovly 2018).

One of the best-studied subfamilies of receptor tyrosine kinases is the ErbB family with its four receptors HER1-4 and a wide range of related growth factors (Schlessinger 2000). All four receptors HER1-4 are structurally related to the epidermal growth factor receptor (EGFR/ErbB1/HER1) which was identified first in 1975 (Carpenter et al. 1975). The discovery of HER2 (Coussens et al. 1985), HER3 (Kraus et al. 1989), and HER4 (Plowman et al. 1993) followed in subsequent years. The ErbB receptor family is involved in a variety of biological activities such as cell differentiation, cell migration, and organ development (Fraguas, Barberán, and Cebrià 2011; Park et al. 2001; Casalini et al. 2004). Mutations in these proteins may lead to dysfunctions resulting in various diseases, such as cancer and inflammation. This renders the ErbB family a prominent therapeutic target (de Bono and Rowinsky 2002; Stern 2000).

EGFR, HER3, and HER4 bind in total eleven known cognate ligands. A first group of ligands, comprising amphiregulin, epidermal growth factor (EGF), epigen, and transforming growth factor α (TGFα), exclusively binds to and activates EGFR. The second group of ligands, comprising betacellulin, epiregulin (EREG), and the ectodomain shredded heparin-binding EGF-like growth factor, bind EGFR as well as HER4. The third group are the neuregulins (NRG) 1-4, which all bind HER4, and NRG1 and NRG2 also bind to HER3 (Macdonald-Obermann and Pike 2014). HER2 is known as an orphan receptor, without any known ligand. Activation occurs through the formation of heterodimers with the other three receptors from this sub-family (Graus-Porta et al. 1997), which is likely facilitated by the extracellular domain of HER2 adopting a conformation similar to that of ligand-bound EGFR, HER3, or HER4 (Cho et al. 2003). EGF and TGFα strongly bind to and activate EGFR, leading to EGFR homodimerization (Schlessinger 2000) and also the formation of EGFR/HER2 heterodimers (Macdonald-Obermann and Pike 2014). EREG binds to HER4 and EGFR, and the formation of heterodimeric complexes with HER2 was reported (Trenker and Jura 2020). NRGβ1 binds to both HER4 and HER3 (Chan et al. 1995; Jones, Akita, and Sliwkowski 1999). It is interesting to note that these four receptors orchestrate many different cellular functions and that this might be regulated by their ligands displaying different receptor binding specificities and affinities, and a different propensity to engage into heterodimers (Sweeney and Carraway 2000).

Single-molecule tracking (SPT) is a powerful method to investigate the mobility of membrane proteins and to associate this information with the activation states of receptors (Momboisse et al. 2022; Ibach et al. 2015; Martínez-Muñoz et al. 2018). For example, a recent study revealed that the receptor tyrosine kinase VEGFR-2 shows diverse modes of activation following the binding of its cognate ligand VEGF (da Rocha-Azevedo et al. 2020).

Here, we used SPT to measure the diffusion of single HER2 receptors in the plasma membrane of living cells in resting and ligand-stimulated conditions. We extracted the diffusion coefficient and mode, and from that inferred the activation of HER2 through heterodimerization with ligand-binding receptors. We observed that the global diffusion coefficient of HER2 decreases in ligand-treated cells. Analysis of the diffusion mode revealed that the fraction of immobile receptors increased at the expense of the freely diffusing population. For EGFR-binding ligands, we observed a stronger activation of HER2 in cells treated with EGF, compared to cells treated with TGFα. Also, we found a stronger activation of HER2 in cells treated with EREG, targeting HER4 and EGFR, as compared to NRGβ1, targeting HER3 and HER4. We further refined the experimental protocol and followed the diffusion coefficient and the different diffusion modes over time for a total of 25 min by measuring multiple cells sequentially. We show that the population of immobile receptors over time correlates with the phosphorylation level of HER2, which was determined at different time points using western blotting. From the time-resolved SPT data, we obtained a temporal profile of HER2 activation in live cells that showed specific features in activation strength, time of maximum activation, and desensitization profile for each ligand.

## Materials and Methods

### Cell culture

HeLa cells (German Collection of Microorganisms and Cell Cultures GmbH, Braunschweig, Germany) were cultured in growth medium (GM), consisting of high glucose Dulbecco’s modified Eagle medium (DMEM) with 1% GlutaMAX, 10% fetal bovine serum (FBS), 100 U/mL penicillin and 100 μg/mL streptomycin (all from Gibco Life Technologies, Waltham, MA, USA), at 37 °C and 5% CO_2_ in an automatic CO_2_ incubator (Model C150, Binder GmbH, Tuttlingen, Germany). For SPT experiments, round coverglasses (25 mm diameter, 0.17 mm thickness, VWR International, Radnor, PA, USA) were passivated and functionalized with PLL-PEG-RGD (Harwardt et al. 2017) and inserted into 6-well plates (Greiner Bio-One, Kremsmünster, Austria). (Harwardt et al. 2017). Cells were seeded to a density of 5 × 10^4^ cells/well for SPT experiments with HER2 and 20 × 10^4^ cells/well for SPT experiments with TMD and GPI and grown at 37 °C and in 5% CO_2_ for 3 days.

### Nanobody expression and purification

The HER2-specific nanobody, 2Rs15d (Vaneycken et al. 2011), was expressed in *E*.*coli* with a C-terminal click handle for site-specific conjugation, as previously described (Teodori et al. 2022). In brief, WK6 *E. coli* cells were co-transformed with a suppressor plasmid, pUltra, and an expression plasmid, pMECS, encoding the nanobody sequence (**Supplemental Note 1**) with an N-terminal pelB leader sequence, a C-terminal hexahistidine tag (6xHis) and an amber stop codon (TAG) positioned on the C-terminal right before the 6xHis-tag. A discrete colony was grown at 37 °C, 220 rpm until optical density (OD600) reached between 0.8 and 0.9. At this OD the unnatural amino acid, 4-azido-L-phenylalanine (1 mM or 0.202 g/L), was added to the culture following induction with IPTG (1 mM) and grown at 18 °C and 200 rpm for a total of 16 h. The next day, cells were harvested and the nanobodies extracted by periplasmic extraction following affinity chromatography purification using Ni-NTA column on an Äkta Start System (Cytiva). The purity of the nanobodies was verified by SDS-PAGE gel and Urea PAGE gel.

### Nanobody labeling

The azide-modified 2Rs15d nanobody was reacted with a 2.5-fold molar excess Cy3B-PEG6-DBCO in 1x PBS in a total volume of 100 μL at 21 °C and 600 rpm for 16 h. SDS-PAGE gel evaluation of the reaction showed a 100% labeling of the nanobody. Excess dye was removed using a PD MiniTrap G-10 gravity column (Cytiva) following the manufacturer’s recommendations. The Cy3B-labeled nanobody was eluted in a total volume of 0.5 mL 1x PBS. The purity of the eluted fractions was assessed by SDS-PAGE gel prior to use for further experiments.

### Sample preparation

Coverglasses were mounted into custom-built holders and rinsed once with 600 μL 1x Live Cell Imaging Solution (LCIS) (Invitrogen, Waltham, MA, USA). 600 μL prewarmed LCIS was added to holders and cooled to room temperature over 15 min. HER2 was labeled with a Cy3B-labeled nanobody (2Rs15d) (Teodori et al. 2022) at a concentration of 2 nM and 10 min prior to the measurements.

For stimulated cells, either 20 nM epidermal growth factor (EGF) (#AF-100-15), transforming growth factor alpha (TGFα) (#100-16A), neuregulin beta 1 (NRGβ1) (#100-03) or epiregulin (EREG) (#100-04) (all from PeproTech, Waltham, MA, USA) were added 5 min after measurement start. SPT experiments were conducted between 21-23 °C (**Figure S9**).

As negative controls, an artificial transmembrane protein (TMD) fused to monomeric enhanced green fluorescent protein (mEGFP) (Wilmes et al. 2020) and a fusion construct of mEos3.2. and glycophosphatidylinositol (GPI)-anchor signal peptide of the human folate receptor (Harwardt et al. 2019) was used. The pSems-mEGFP-TMD plasmid was kindly provided by Jacob Piehler (University of Osnabrück, Germany). 100 ng/well pSems-mEGFP-TMD plasmid and 2.25 μg/well sheared salmon sperm DNA (#AM9680, Invitrogen) or 500 ng/well of the pN1-GPI-mEos3.2. plasmid and 1.5 μg/well sheared salmon sperm DNA were transfected using Lipofectamin 3000 (Thermo Fisher Scientific) following the manufacturer’s protocol in 6-well plates. Transfected cells were incubated overnight at 37 °C and 5% CO_2_. Prior to microscopy experiments, cells were washed with 600 μL prewarmed LCIS, 600 μL fresh LCIS added, and incubated for 15 min at room temperature. The mEGFP-TMD was eventually labeled with 0.5 nM of the mEGFP-targeting FluoTag®-Q nanobody labeled with AbberiorStar635P (NanoTag Biotechnologies, Göttingen, Germany) next to EGF addition for the ligand stimulated condition 5 min prior to measurements. For pN1-GPI-mEos3.2, the fluorescent protein mEos3.2. was tracked, and EGF was added for the ligand-stimulated condition 5 min prior to measurements.

### Single-molecule microscopy

Data were acquired on a commercial widefield microscope (N-STORM; Nikon, Düsseldorf, Germany) equipped with an oil-immersion objective (100x Apo TIRF oil; NA 1.49), operated in total internal reflection fluorescence (TIRF) mode, and a 1.5x magnification lens inserted into the detection beam path for measurements of HER2 and GPI. As an excitation light source for Cy3B and mEos3.2, a laser emitting at 561 nm was used and operated at an irradiation intensity of 6.3 W/cm^2^ or 30 W/cm^2^ respectively. The fluorescent protein mEos3.2 was photoconverted to its orange fluorescent state using a 405 nm laser with the intensity adapted to the expression level in single cells. AbberiorStar635P was excited with a laser emitting at 647 nm at 0.6 kW/cm^2^. Fluorescence emission was detected with an electron-multiplying charge-coupled device (EMCCD; Andor iXon, DU-897U-CS0-BV; Andor, Belfast, UK) using an EM gain of 300 (for Cy3B) or 200 (for AbberiorStar635P and mEos3.2), a pre-amplifier gain of 3 and a read-out rate of 17 MHz with activated frame transfer. Images of 256 × 256 pixels were acquired with 157 nm pixel size for experiments with TMD and 105 nm pixel size for experiments with HER2 and GPI.

The microscope was controlled by *NIS Elements* (v4.30.02, Nikon) and *μManager (Edelstein et al. 2014)*. For each cell, a total of 1000 frames were recorded at an integration time of 20 ms. To record a time series, 25 cells were imaged sequentially during an acquisition time of about 30 minutes. To record time series for cells that were stimulated with a ligand, the ligand was added after 5 cells were measured.

### Data analysis

Single emitters were localized with *ThunderSTORM* version dev-2016-09-10-b1 (Ovesný et al. 2014) using a plugin for Fiji (Schindelin et al. 2012). Parameters for the tracking analysis (*precision, exp_noise_rate, diffraction_limit, exp_displacement*, and *p_bleach*) were determined from localization data following a previously published procedure using *SPTAnalyser* (https://github.com/JohannaRahm/SPTAnalyser) (J. V. Rahm et al. 2021; J. Rahm et al. 2022). The *exp_noise_rate* and *precision* were calculated individually per cell whereas the parameters *diffraction_limit* (HER2: 17 nm, TMD: 23 nm, GPI: 30 nm), *exp_displacement* (HER2: 117 nm, 82 nm, TMD: GPI: 165), and *p_bleach* (HER2: 0.064, TMD: 0.0138, GPI: 0.095) were averaged and used globally for all cells. The switching probability was set to 0.01. Localizations were connected to trajectories using the software package *swift* (v0.4.2) (Endesfelder et al., manuscript in prep.) with the aforementioned parameters. MSD analysis (fitting length of 4 data points for calculation of diffusion coefficients), filtering (minimal trajectory length of 20), and assignment of diffusion types were performed in *SPTAnalyser* (J. V. Rahm et al. 2021). Segments were classified as immobile with a threshold of a minimal diffusion coefficient (D_min_(HER2) = 0.0084 μm^2^ s^-1^, D_min_(TMD) = 0.0037 μm^2^ s^-1^, D_min_(GPI) = 0.0086 μm^2^ s^-1^) calculated from the third quartile of the dynamic localization precision (Michalet 2010; Rossier et al. 2012). Diffusion coefficients and modes were calculated per individual cell and averaged over all cells. For comparison of global values for diffusion coefficients and fractions of diffusion types, the data recorded for cells from time intervals of 0 to 20 min were grouped. To correct for fluctuations between the resting condition in different measurements, the relative occurrence of diffusion modes in the first five minutes of measurements (−5 min interval, prior to ligand addition) were aligned (uncorrected data are shown in **Figure S6**). Time course analyses were performed by grouping diffusion coefficients and modes in time groups of 1 or 5 min to minimize the contribution of cell heterogeneity (da Rocha-Azevedo et al. 2020).

*Swift* version 0.4.2, used in this manuscript, and all subsequent versions of the *swift* software, as well as documentation and test data sets, can be obtained on the *swift* beta-testing repository (http://bit.ly/swifttracking). The home-written software *SPTAnalyser* in Python (3.7.6) estimates parameters for tracking with *swift* and executes diffusion state analysis and transition counting. *SPTAnalyser* has a graphical user interface with adaptable analysis parameters and assists in processing large amounts of data by creating macros for *ThunderSTORM* and batch files for *swift. SPTAnalyser* is compatible with *PALMTracer* (Bordeaux Imaging Center), which is software for localization and tracking available as a plugin for *MetaMorph* (Molecular Devices, Sunnyvale, CA, USA). The source code of *SPTAnalyser*, together with a detailed manual, is available from https://github.com/JohannaRahm/SPTAnalyser.

### Western blotting

0.9 × 10^6^ HeLa cells were seeded onto 10 cm cell culture dishes (Greiner Bio-One, Kremsmünster, Austria) in GM and grown at 37 °C and 5% CO_2_. In the evening of the third day, cells were starved with serum-free GM overnight. Cells were stimulated with 20 nM of one of the respective ligands EGF, TGFα, NRGβ1, or EREG in serum-free GM and incubated for 2, 5, or 30 min. For the control western blot, cells were incubated with 20 nM EGF and 2 nM nanobody for 5 min. Afterwards, cells were rinsed with ice-cold 1x Dulbecco’s phosphate-buffered saline (PBS) pH 7.4 (Gibco Life Technologies, #14040133) and incubated for at least 2 min on ice prior to adding lysis buffer consisting of 150 mM NaCl, 50 mM Tris-HCl pH 7.4, 10 mM NaF, 1 mM Na_3_VO_4_, 1 mM EDTA, 1%(v/v) Triton X-100, 0.5%(w/v) Na-deoxycholate, 0.1%(w/v) SDS (all from Sigma-Aldrich, St. Louis, MO, USA), and ¼ of a cOmplete Mini EDTA-free protease inhibitor tablet (Roche, Basel, Switzerland) in 10 mL buffer. Cells were scraped and the collected lysate was shaken at 4 °C for 5 min at 750 rpm (Thermo-Shaker, Universal Labortechnik GmbH & Co. KG, Leipzig, Germany). Lysate and cell fragments were separated by centrifugation at 4 °C for 20 min at 12,000 rpm (Centrifuge 5418 R, Eppendorf, Hamburg, Germany). Protein concentrations in the supernatant were determined using the Pierce Micro BCA Protein Assay kit (Thermo Fisher Scientific, Waltham, MA, USA) according to the manufacturer’s protocol.

SDS-PAGE was performed to analyze the time-dependent phosphorylation of HER2 upon ligand stimulation. Precast 4-20% gradient SDS-PAGE gels (Mini-PROTEAN^®^ TGX™, BioRad Laboratories, Hercules, CA, USA) were mounted in a cask filled with running buffer (25 mM Tris base, 190 mM glycine, 3.5 mM SDS, pH 8.3, all from Sigma-Aldrich). 50 μg protein were prepared in 20%(v/v) loading dye (250 mM Tris-HCl (pH 6.8), 8%(w/v) SDS, 0.1%(w/v) bromophenol blue (all from Sigma-Aldrich), and 40%(v/v) glycerol (Carl Roth, Karlsruhe, Germany)), supplied with 0.1 M dithiothreitol (Sigma-Aldrich), heated to 95 °C for 5 min and loaded onto the gel with PageRuler™ Prestained Protein Ladder (Thermo Fisher Scientific) as a reference marker. Gels were run at 60 V for 10 min to allow the samples to enter the gel and then at 200 V for 45 min.

Gels were blotted for 7 min using an iBlot Gel Transfer Device (Invitrogen). All further incubation steps were performed under agitation at room temperature if not stated otherwise. First, blots were incubated in blocking buffer (5%(w/v) milk powder (nonfat dry milk, Cell Signaling Technology, Danvers, MA, USA)) in TBST containing 25 mM Tris base, 150 mM NaCl, and 0.05%(v/v) Tween-20 (all from Sigma-Aldrich) in water, pH 7.6, for 1 h. After washing three times with TBST for 5 min, the blots were incubated with primary antibodies against HER2 (rabbit anti-HER2 (Y1221/1222), Cell Signaling Technology #2243, diluted 1:500 for EGF-, TGFα- and EREG-stimulated samples, diluted 1:200 for NRGβ1-stimulated samples) and a housekeeping gene (rabbit anti-actin, abcam #ab14130, diluted 1:40,000 for all conditions) in TBST supplemented with 5%(w/v) BSA (Sigma-Aldrich) at 4 °C overnight. Blots were washed three times with TBST for 5 min prior to the addition of the secondary antibody (goat anti-rabbit tagged with horseradish peroxidase, Jackson ImmunoResearch, West Grove, PA, USA, #111-035-003, diluted 1:20,000) in TBST supplemented with 5%(w/v) BSA (Sigma-Aldrich). The secondary antibody was incubated for 3 h. Afterwards, blots were washed four times with TBST for 5 min, 10 min, 15 min, and 15 min, respectively. Lastly, washing with TBS was performed for 5 min. For imaging, blots were treated with SuperSignal West Femto Maximum Sensitivity Substrate (Thermo Fisher Scientific) according to the manufacturer’s protocol, and bands were detected on a CHEMI-only chemiluminescence imaging system (VWR, Radnor, PA, USA).

### Statistical analysis

Statistical analysis was performed with OriginPro 2022 (v9.9.0.225, OriginLab Corporation, Northampton, MA, USA). Mean values were calculated for all diffusion properties of individual cells and displayed with their respective standard errors. Populations were tested for being normally distributed using the Shapiro-Wilk test (α = 0.05). As some populations rejected this hypothesis, non-parametric tests were chosen for comparing data. The Mann-Whitney-U test was used to compare distributions from different treatment groups whereas Wilcoxon signed-rank tests were used to validate data from the same treatment group. The following classification of significance levels was used: p ≥ 0.05 no significant difference (not labeled), p < 0.05 significant difference (*), p < 0.01 very significant difference (**), p < 0.001 highly significant difference (***).

## Results

We investigated the activation strength of HER2 in live HeLa cells treated with different ligands that target its heterodimerization partners EGFR, HER3, or HER4. For that purpose, we measured the diffusion coefficient and type of HER2 in unstimulated and ligand-stimulated HeLa cells using single-particle tracking (SPT) (Shen et al. 2017) with the fluorophore-conjugated nanobody 2Rs15d (Vaneycken et al. 2011) (**Figure 1A**). This nanobody was found to only bind domain I of the HER2 receptor (D’Huyvetter et al. 2017) and to not compete with HER2-specific inhibitors trastuzumab and pertuzumab (Vaneycken et al. 2011), which target the dimerization interface of the receptor (Leahy 2004; Cho et al. 2003; Franklin et al. 2004). Hence, this nanobody does not, or only to a small extent, impair the formation of heterodimers of HER2 with EGFR, HER3, and HER4 which occurs through domain II of the HER2 receptor (Hu et al. 2015). We were able to confirm this by western blot analysis (**Figure S1**).

**Figure 1.**
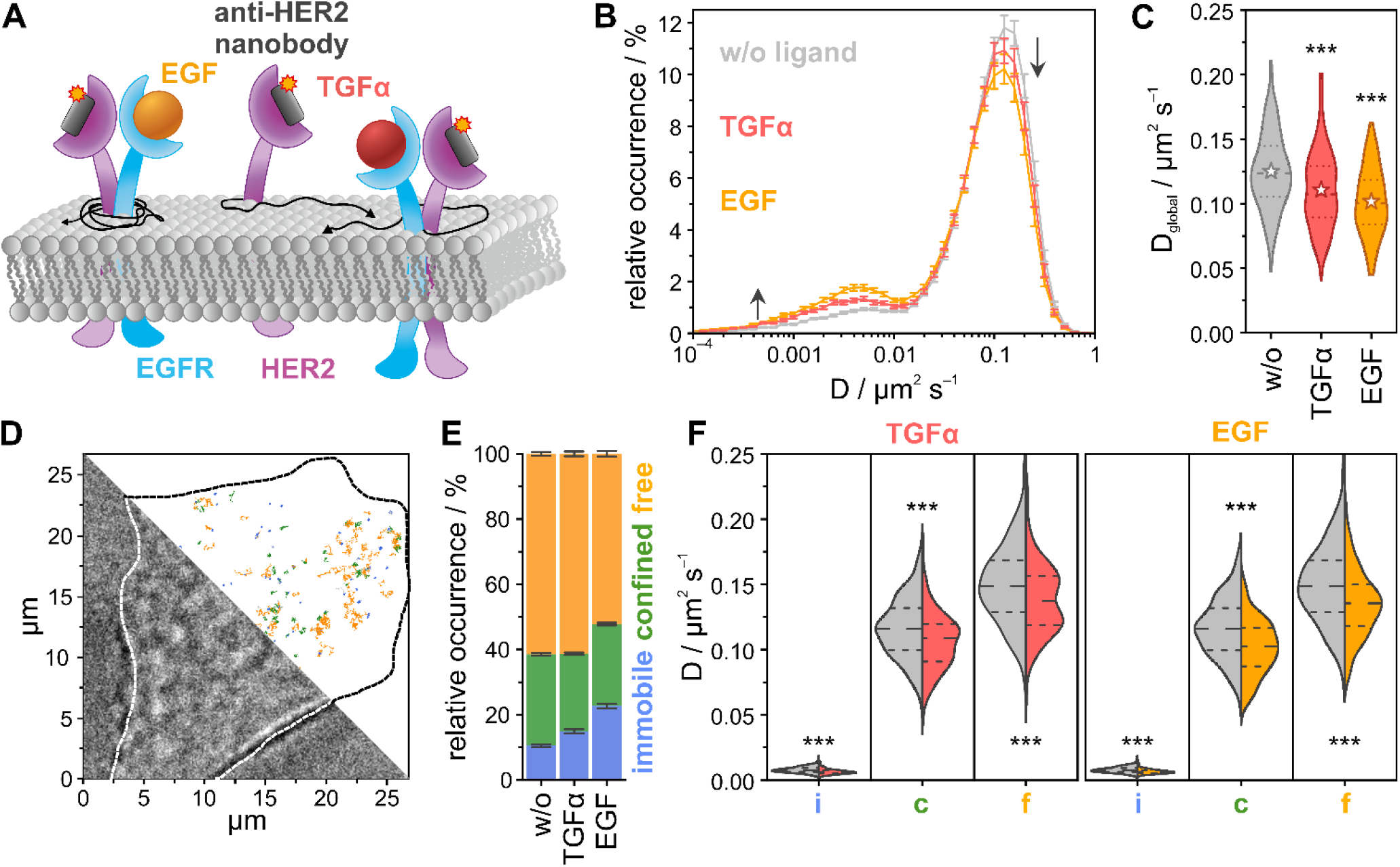
Single-particle tracking of HER2 in live HeLa cells treated with the EGFR-targeting ligands EGF and TGFα. (A) HER2 was targeted with a Cy3B-labeled anti-HER2 nanobody, and the mobility of HER2 was measured in the absence and presence of EGF and TGFα. (B) Distribution of diffusion coefficients for resting (gray), EGF- (orange), and TGFα-treated (red) HeLa cells at 22 °C. (C) Global diffusion coefficient per condition (violin plots with dotted lines marking the quartiles, dashed lines the median, and stars representing mean values). (D) Exemplary bright-field image of a living HeLa cell treated with EGF, and single-molecule trajectories colored for their diffusion mode, immobile (blue), confined (green), free (orange). (E) Relative occurrences of immobile, confined, and freely diffusing HER2 receptors in live HeLa cells. (F) The diffusion coefficient for the individual diffusion modes immobile (i), confined (c), and free (f) (violin plots, dense dashed lines represent the quartiles, loosely dashed lines represent the median). The data shown was assembled from 160 cells. Error bars are defined by SEMs; p > 0.05 no significant difference (no label), p < 0.05 significant difference (*), p < 0.01 very significant difference (**), p < 0.001 highly significant difference (***).

First, we investigated how ligand activation of EGFR, a heterodimerization partner of HER2 (Graus-Porta et al. 1997; Gulliford et al. 1997), impacts the mobility of HER2 receptors. We selected the ligands EGF and TGFα, which exclusively bind to EGFR (Olayioye et al. 2000), and measured the diffusion coefficient of single HER2 receptors in the basal plasma membrane of live HeLa cells. In untreated cells, we found a bimodal distribution of the diffusion coefficient of HER2 (**Figure 1B**), from which we calculated a global diffusion coefficient of D_global_ = 0.125 ± 0.005 μm^2^ s^−1^ (**Figure 1C**). In cells treated with EGFR-targeting ligands EGF or TGFα, we found a decrease in the fraction of HER2 receptors with high diffusion coefficients, and an increase in the fraction with low diffusion coefficients, with EGF showing a stronger effect than TGFα (**Figure 1B**). This was mirrored in a decrease of the global diffusion coefficients, which were calculated to D_global,EGF_ = 0.102 ± 0.005 μm^2^ s^−1^ and D_global,TGFα_ = 0.111 ± 0.005 μm^2^ s^−1^ for cells treated with EGF or TGFα, respectively (**Figure 1C**). Next, we analyzed the mode of diffusion (J. V. Rahm et al. 2021), and distinguished immobile HER2 receptors from those showing confined or free diffusion (**Figure 1D-F**). For HER2 in untreated cells, we found 10.5 ± 0.4% immobile receptors, whereas 28.1 ± 0.4% and 61.4 ± 0.6% showed confined or free diffusion, respectively (**Figure 1E**). In cells treated with EGF and TGFα, we observed an increase in the immobile fraction of HER2 receptors, at the expense of freely diffusing receptors. For cells treated with EGF, we determined the fraction of immobile HER2 receptors to 22.7%, which corresponds to an increase of 116%. In cells treated with TGFα, we found a smaller increase in the immobile fraction to 14.9%, which corresponds to an increase of 42%. At the same time, we found that the diffusion coefficient of the fraction of freely or confined diffusing HER2 receptors was reduced in cells treated with EGF or TGFα (**Figure 1F**). Changes in the quantity of immobile receptors indicate HER2 activation and ligand-specific responses. The amount of immobile receptors in untreated cells could partly also arise from unspecifically bound nanobody to the cell surface. As a control experiment, we measured the diffusion coefficient and type of a transmembrane domain (TMD) peptide conjugated to mEGFP (mEGFP-TMD) that was targeted with a fluorophore-labeled anti-GFP nanobody (Wilmes et al. 2020) and of a GPI conjugated to mEos3.2 (Harwardt et al. 2018). We found no significant difference in the diffusion coefficient of, nor changes in the diffusion type for mEGFP-TMD or GPI-mEos3.2 in untreated and EGF-treated cells (**Figure S2**; **Tables S1-S8**). Furthermore, we monitored key parameters of all SPT experiments in untreated and ligand-treated cells and calculated the average number of trajectories and segments per cell and trajectory as well as segment lengths for all conditions (**Figure S3**).

Next, we investigated the activation of HER2 in cells treated with EREG, which predominantly binds HER4 as well as EGFR, and with NRGβ1, which binds to HER3 and HER4 (Olayioye et al. 2000), by measuring the mobility of HER2 receptors in live HeLa cells (**Figure 2A**). We found that the bimodal distribution of the diffusion coefficient of HER2 showed small changes in cells treated with EREG or NRGβ1, as compared to untreated cells (**Figure 2B**). This is reflected in smaller changes of the global diffusion coefficient, D_global,EREG_ = 0.113 ± 0.005 μm^2^ s^−1^ and D_global,NRGβ1_ = 0.121 ± 0.005 μm^2^ s^−1^, for EREG and NRGβ1, respectively (**Figure 2C**). The analysis of diffusion types of single trajectories (**Figure 2D**) showed that the fraction of immobile receptors increased from 10.5 ± 0.4% in untreated cells to 12.5 ± 0.5% and 16.5 ± 0.5% in cells treated with NRGβ1 or EREG, respectively, and at the expense of a decrease of the mobile fraction (**Figure 2E**). Again, we found that the diffusion coefficient of the fraction of freely or confined diffusing HER2 receptors was reduced in cells treated with EREG or NRGβ1 (**Figure 2F**).

**Figure 2.**
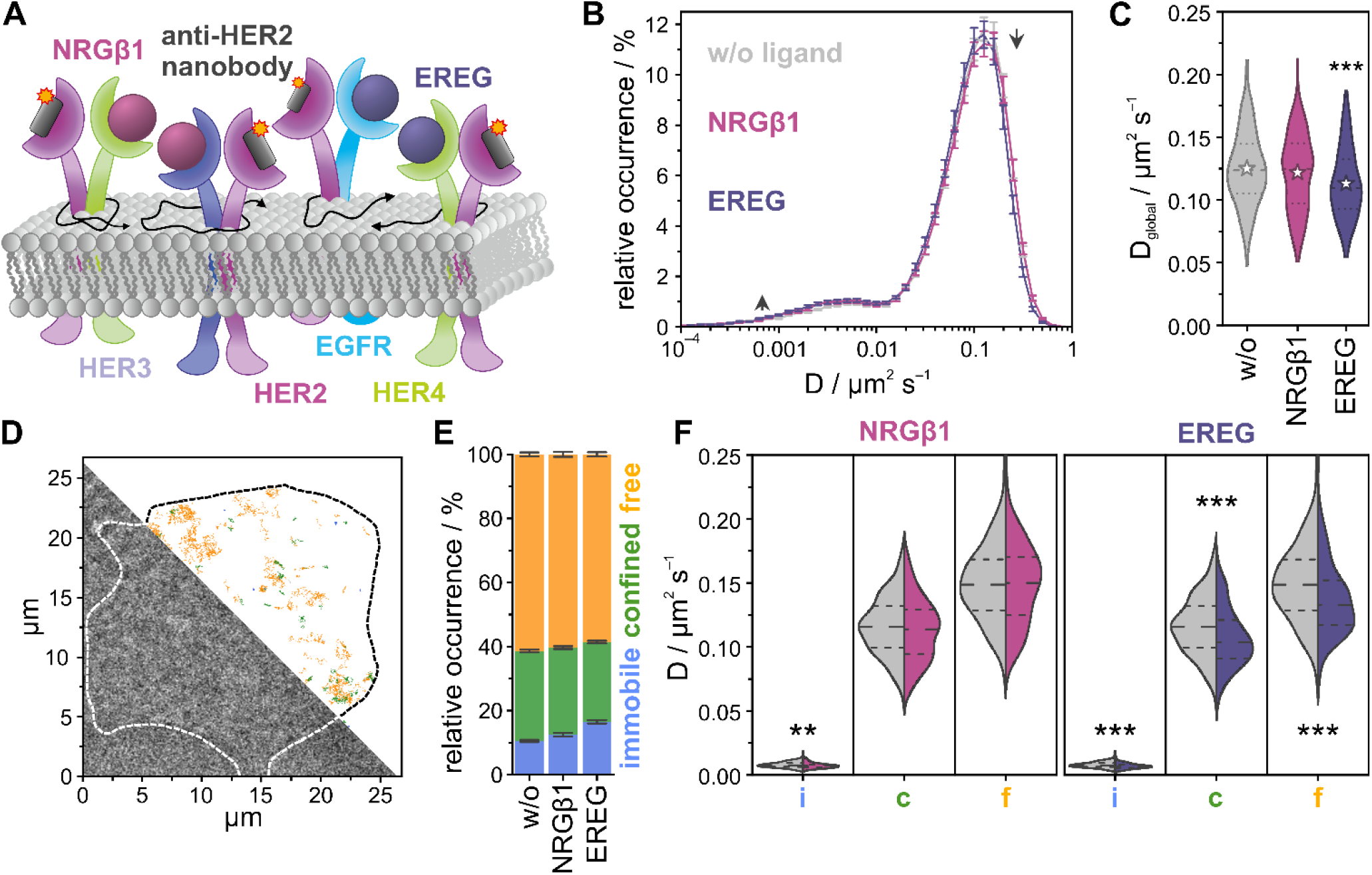
Single-particle tracking of HER2 in live HeLa cells treated with the ligands EREG and NRGβ1. (A) HER2 was targeted with a Cy3B-labeled anti-HER2 nanobody, and the mobility of HER2 was measured in the absence and presence of EREG and NRGβ1. (B) Distribution of diffusion coefficients for resting (gray), EREG- (purple), and NRGβ1-treated (lilac) HeLa cells at 22 °C. (C) Global diffusion coefficient per condition (violin plots with dotted lines marking the quartiles, dashed lines the median, and stars representing mean values). (D) Exemplary bright-field image of a living HeLa cell treated with NRGβ1, and single-molecule trajectories colored for their diffusion mode, immobile (blue), confined (green), free (orange). (E) Relative occurrences of immobile, confined, and freely diffusing HER2 receptors in live HeLa cells. (F) The diffusion coefficient for the individual diffusion modes immobile (i), confined (c), and free (f) (violin plots, dense dashed lines represent the quartiles, loosely dashed lines represent the median). The data shown was assembled from 160 cells. Error bars are defined by SEMs; p > 0.05 no significant difference (no label), p < 0.05 significant difference (*), p < 0.01 very significant difference (**), p < 0.001 highly significant difference (***).

The experiments were further refined to follow the activation of HER2 receptors in live cells over longer time periods, using the diffusion coefficient and type as proxies for the formation of heterodimers with EGFR, HER3, or HER4. Since the observation time of HER2 receptors bound to a fluorophore-labeled nanobody is limited by photobleaching, the time window for the observation of a single cell is too short to follow changes related to signaling activation that typically occur at the time scale of minutes (Kiso-Farnè and Tsuruyama 2022). To bypass this limitation, we established an experimental procedure in which we measured many cells sequentially, giving each cell its own time stamp (see Methods) (**Figure 3A**). Using that procedure, we measured the diffusion coefficient and type of single HER2 receptors over a period of 25 min in the same well. The respective ligand was added after 5 min, to record reference data for untreated cells; this enabled it to follow ligand-specific changes in diffusion coefficient and type for a time of 20 min.

**Figure 3.**
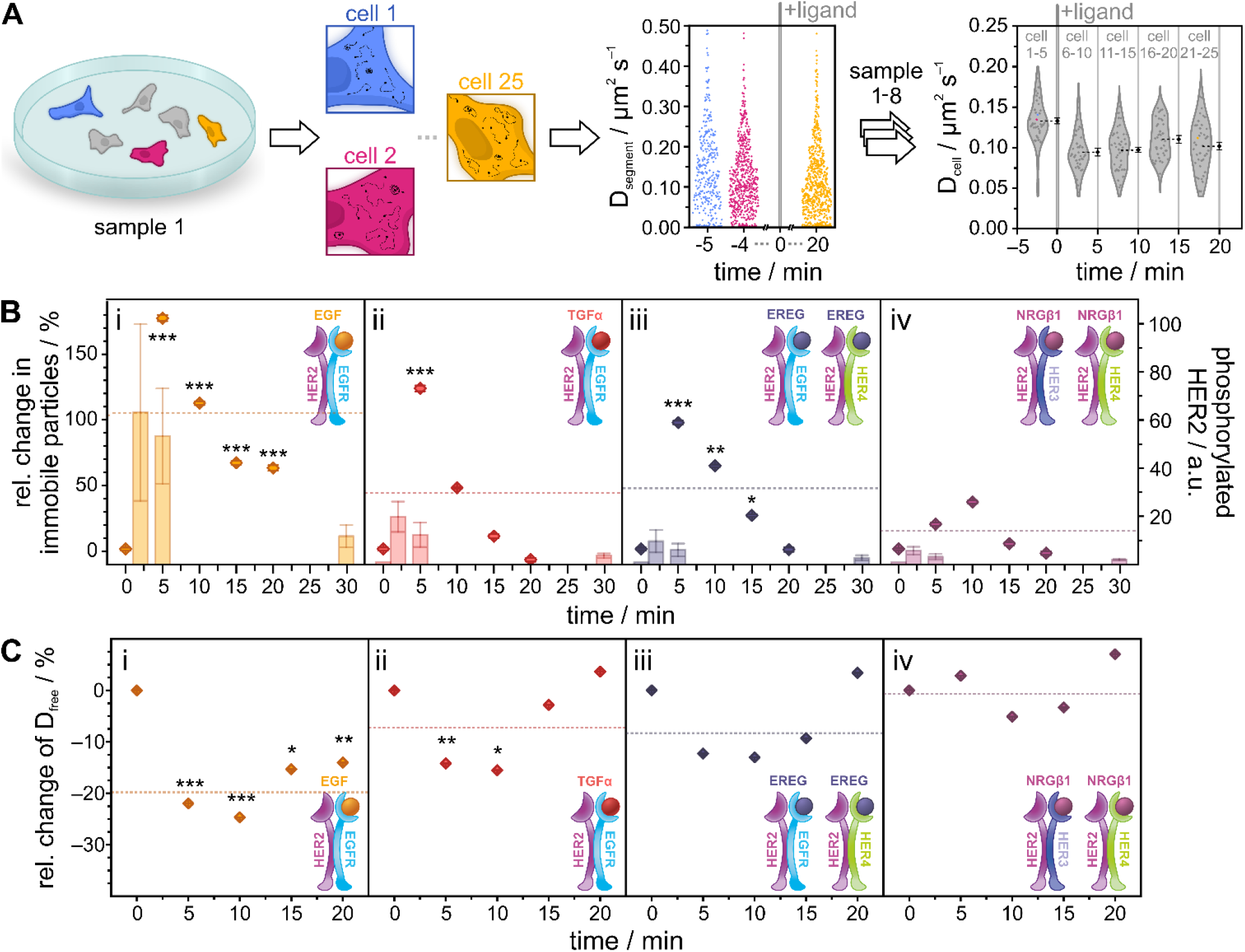
Temporal response of HER2 activation in living HeLa cells following treatment with EGF, TGFα, EREG, or NRGβ1. (A) Schematic representation of the time-course SPT experiment. Cells were seeded sparsely and imaged sequentially. After imaging 5 cells in resting condition, the respective ligand was added to the cell dish and the measurement continued. Diffusion mode and coefficients were calculated per cell and pooled into 5 min time intervals. Diamonds represent mean diffusion coefficients per segment (colored) or cell (grey; mean values are colored in black with error bars representing the SEM). (B) Relative change in the fraction of immobile particles (dot plots) plotted against time. Bars show HER2 phosphorylation obtained from western blots (N = 3). (C) Relative change in the diffusion coefficient of freely diffusing particles over time. Relative changes were calculated from mean values of 40 cells per interval. Receptor models indicate the expected ligand-orchestrated interactions between HER2 and other receptors of the family. The dotted lines represent mean values of the relative change over the time of ligand stimulation. Error bars in dot plots represent the standard error of the difference (SED); error bars in bar plots show the standard error of the mean (SEM). Significance was tested for stimulated cells vs. untreated cells from the same sample before calculating the relative change; p > 0.05 no significant difference (no label), p < 0.05 significant difference (*), p < 0.01 very significant difference (**), p < 0.001 highly significant difference (***).

We analyzed the diffusion type and coefficient over time for all four ligands, EGF, TGFα, EREG, and NRGβ1 (**Figure 3B**,**C; S4-S7**). Considering our previous result of an increase in the immobile fraction of HER2 in ligand-treated cells (**Figures 1E; 2E**), we followed the population of the immobile fraction over time (**Figures 3B; S6C**). For all ligands, we found a strong increase of the immobile fraction after 5 min, which represented at the same time the maximum in cells treated with the ligands EGF, TGFα, and EREG. In cells treated with NRGβ1, we found the maximum population of the immobile fraction shifted to ∼10 min. In cells treated with EGF, the fraction of immobile HER2 increased by 180% after 5 min of stimulation before slowly decreasing to ∼60% within the following 15 min (**Figure 3Bi**). In cells treated with TGFα and EREG, the fraction of immobile particles increased to a maximum of ∼120% and 100%, respectively, compared to unstimulated cells (**Figures 3Bii; 3Biii**), while changes observed for cells treated with NRGβ1 were smaller (**Figure 3Biv**). While the population of immobile particles returned to the level found in untreated cells for TGFα, EREG, and NRGβ1, this was not found for EGF. To correlate the increase in the population of the immobile state with the activation of HER2, we performed a western blot analysis of phosphorylated HER2 for all four ligands at different time points (**Figure 3B; S8**). We found the maximum population of phosphorylated HER2 around 2-5 min for all four ligands, similar to the population maxima of the immobile state. The activation strengths for phosphorylated HER2 were strongest for EGF, followed by TGFα and EREG (**Figure 3B; S8**).

The population of the fractions of freely and confined diffusing HER2 also showed a first response after 5 min, with different temporal signatures and strengths for the different ligands (**Figures S4; S6A**,**B**). For all four ligands, we measured a reduced diffusion coefficient for the freely diffusing population of HER2, amounting to ∼5% (NRGβ1), ∼15% (TGFα, EREG), and ∼25% (EGF) (**Figures 3C; S7A**). We further found that this change was shifted to later time points of ∼10 min for all receptors. For cells treated with TGFα, EREG, or NRGβ1, we found that after 20 min, the diffusion coefficient showed similar values as in untreated cells, whereas this was not the case for cells treated with EGF. The diffusion coefficient of confined HER2 receptors shows a similar temporal signature and strength (**Figures S5A; S7B**), while smaller effects were found for the diffusion coefficient of immobile HER2 (**Figures S5B; S7C**).

## Discussion

Using live-cell single-particle tracking and data analysis, we extract the diffusion coefficients and modes of HER2 in the native plasma membrane of living HeLa cells. We found that HER2 molecules in the plasma membrane of unstimulated cells exhibit heterogeneity in mobility, including free and confined diffusing as well as immobile receptors. In order to attribute these states to their potential activity, we first monitored how the population of the mobility states changes upon ligand treatment of known heterodimerization partners of HER2. For all four ligands investigated, EGF, TGFα, EREG, and NRGβ1, we find a decrease in free diffusing HER2 and an increase in immobile HER2. This indicates that free diffusing HER2 promotes encounters with the interaction partners EGFR, HER3, and HER4 and that the heterodimers enrich into immobile receptor complexes. A similar observation was reported for EGFR, which populates a slow diffusion state upon binding EGF (Chung et al. 2010). In order to further support this interpretation for HER2, we performed western blotting and found an increase in phosphorylated HER2 that correlated with the increase of immobile particles. We also found that the global diffusion coefficient of HER2 derived from all HER2 molecules without grouping into diffusion modes was reduced in cells treated with EGF, TGFα, or EREG, with the response being strongest for EGF. This reduction was also reflected in the diffusion coefficient for free and confined diffusing HER2 molecules in cells treated with EGF, TGFα, or EREG.

Our single-molecule imaging method allowed following the movement of a single HER2 molecule for up to a few seconds, limited by photobleaching. Signaling initiation of ErbB receptors, however, occurs at the time scale of minutes (Hass et al. 2017). An elegant strategy to bridge these two time windows is to measure the mobility of single receptors in many different cells of the same dish sequentially, as it was reported for VEGFR-2 recently and allowed to follow receptor activation after ligand stimulation (da Rocha-Azevedo et al. 2020). We adapted this concept and established a time-course single-particle experiment by measuring HER2 mobility in many cells from the same dish sequentially. We measured the activation of HER2 in response to different ligands that target its heterodimerization partners for up to 30 minutes. Each measured cell was time-stamped, resulting in a temporal profile of HER2 mobility following ligand treatment of cells. This information-rich data informed on the temporal signature of HER2 activation, its strength, and desensitization over time, for the respective ligand. Commonly for all four ligands investigated, we found an enrichment in immobile HER2 molecules peaking at 5-10 minutes after ligand treatment, paralleled by a decrease of free diffusing HER2 molecules. The strength of HER2 activation, measured as the increase of enrichment of immobile particles and supported by western blot data of HER2 phosphorylation (**Figure 4A**), was highest for EGF, followed by TGFα and EREG, and the weakest for NRGβ1. The diffusion coefficient of mobile HER2 molecules decreased for all ligands by 10-25%, with the extent of decrease being highest for EGF, followed by TGFα, EREG, and NRGβ1. These ligand-induced changes in diffusion mode and coefficient are very similar to recently reported results for the receptor tyrosine kinase VEGFR-2 (da Rocha-Azevedo et al. 2020). The temporal profile of HER2 activation by EGF is also in line with a reported systems biology model (Hass et al. 2017).

**Figure 4.**
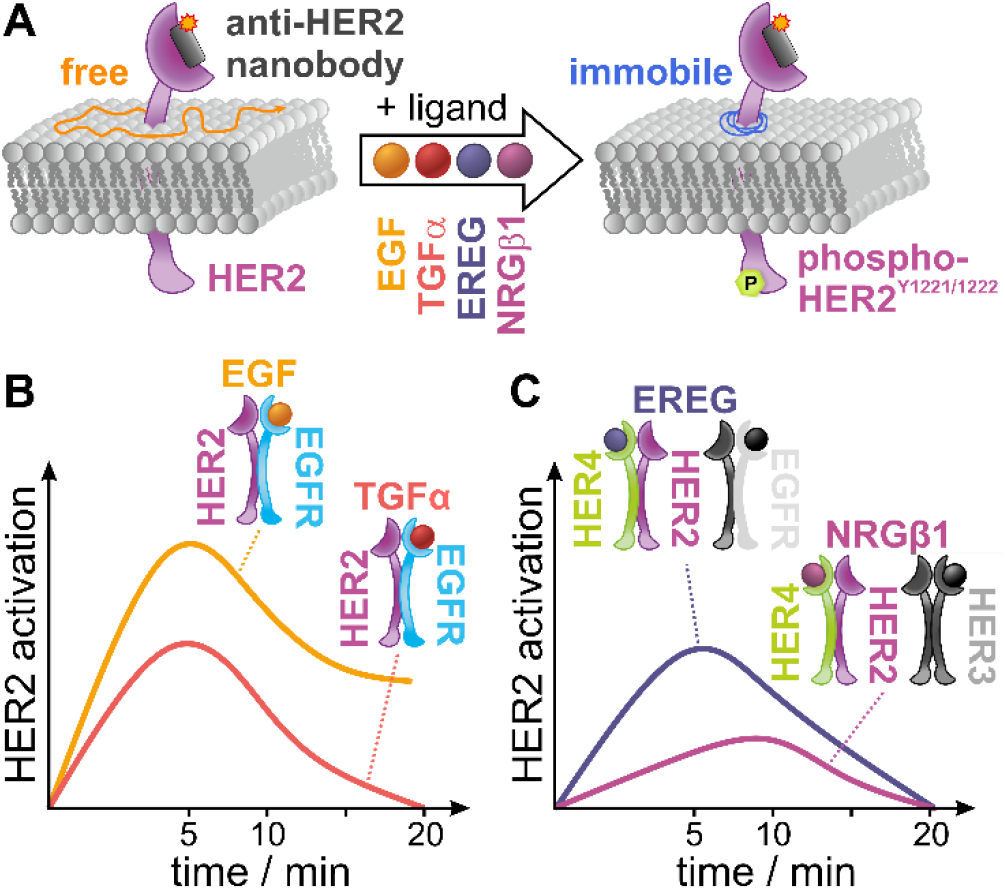
Proposed model for ligand-induced activation of HER2 derived from single-particle tracking data shown in a model-like fashion by means of the active population. (A) In cells treated with EGF, TGFα, EREG, or NRGβ1, the population of immobile HER2 increases at the cost of freely diffusing HER2. This increase scales with the formation of phosphorylated HER2. (B) Temporal profile, strength, and decay of HER2 activation in cells treated with the EGFR-targeting ligands EGF and TGFα. (C) Temporal profile, strength, and decay of HER2 activation in cells treated with the ligand EREG and NRGβ1.

EGF and TGFα bind exclusively to EGFR and with similar affinity (Massagué 1983; Schreiber, Winkler, and Derynck 1986; Lax et al. 1988), yet initiate differential intracellular signaling responses (Ebner and Derynck 1991; Schreiber, Winkler, and Derynck 1986; Korc, Haussler, and Trookman 1987; Barrandon and Green 1987; Wilson et al. 2009, 2012; Knudsen et al. 2014). Both ligands were reported to exhibit an increased affinity for EGFR/HER2 heterodimers compared with EGFR homodimers (Macdonald-Obermann and Pike 2014). The temporal profiles of HER2 activation, derived from an increase in immobile HER2 molecules and the parallel decrease of free HER2 molecules, peak at 5 min for both EGF and TGFα both, but differ in strength, with EGF superseding TGFα (**Figure 4B**). The desensitization of TGFα is complete after 20 min, when both the immobile and free HER2 molecules return to levels in unstimulated cells, yet not for EGF, where about 40% of HER2 molecules remain immobile after 20 min. These results suggest that HER2 is preferably recruited to EGF-bound EGFR, which agrees with biochemical data reporting that TGFα has a reduced ability to recruit EGFR into heterodimers with EGFR (Gulliford et al. 1997). The results also show that the different signaling responses initiated by EGF and TGFα are mirrored in the temporal profile of diffusion mode population and diffusion coefficient.

EREG and NRGβ1 both bind to HER4, with EREG showing weak affinity to EGFR and NRGβ1 also binding to HER3 (Jones, Akita, and Sliwkowski 1999; Riese and Cullum 2014). The temporal profiles of HER2 activation, derived from an increase in immobile HER2 molecules and the parallel decrease of free HER2 molecules, showed a peak at 5 min for EREG and at 10 min for NRGβ1 (**Figure 4C**). In addition, we found a strong activation of HER2 by EREG and a rather weak activation by NRGβ1. NRGβ1 is reported to strongly bind both HER3 and HER4 (Jones, Akita, and Sliwkowski 1999). However, the expression level of HER3 in HeLa cells is lower than that of HER4 (**Figure S9**), and HER3 is reported to mainly localize intracellularly (Chen B, Mao R, Wang H, She J. 2010) (**Supplemental Note 2**), suggesting that the observed activation of HER2 in response to NRGβ1 can be mainly attributed to HER2/HER4 heterodimer formation. EREG is a low-affinity ligand to EGFR (Freed et al. 2017) while binding HER4 with high affinity (Jones, Akita, and Sliwkowski 1999). This suggests that the observed activation of HER2 in response to EREG can be mainly ascribed to HER2/HER4 heterodimer formation. In consequence, the activation strength of HER2 through the formation of HER2/HER4 heterodimers is stronger for EREG than for NRGβ1, indicating a bias in signaling activation.

In summary, we found activation patterns for HER2 in live cells that differed for the four ligands investigated. EGF and TGFα, both binding EGFR, show a stronger activation of HER2 than EREG and NRGβ1, predominantly binding to HER4 in HeLa cells (**Supplemental Note 2**). In part, this might be related to differences in the expression level of EGFR and HER4 (**Figure S9B**), which to some degree influences the probability of encounter of receptors and heterodimer formation. For both pairs of ligands that lead to the formation of the respective heterodimers EGFR/HER2 (binding of EGF or TGFα to EGFR) and HER4/HER2 (binding of EREG or NRGβ1 to HER4), we find a different activation strength of HER2. Since these ligands also initiate differential intracellular signaling responses for their target receptors EGFR (Ebner and Derynck 1991; Schreiber, Winkler, and Derynck 1986; Korc, Haussler, and Trookman 1987; Barrandon and Green 1987; Wilson et al. 2012, 2009; Knudsen et al. 2014) and HER4 (Jones, Akita, and Sliwkowski 1999), this indicates that HER2 heterodimers with EGFR and HER4 show a selective response to the respective ligand or biased signaling.

## Conclusion

We measured the plasma membrane mobility of HER2 in live HeLa cells treated with various ligands targeting the heterodimerization partners EGFR, HER3, and HER4. We extracted diffusion coefficient and type, monitored specific changes in ligand-treated cells, and found different activation strengths for the heteromeric receptor complexes with EGFR and HER3/HER4. By measuring the diffusion properties of single HER2 receptors in many single cells sequentially, we were able to monitor how the diffusion states of HER2 changed over longer time periods. The temporal profile of diffusion states hereby correlated well to the reported kinetics of signaling activation through HER2, indicating that diffusion properties can serve as proxies to follow the activation of HER2 in heteromeric receptor complexes. This allowed us to characterize ligand-specific activation profiles related to the formation of the different heteromeric receptor complexes, for which we found distinguishable activation kinetics, activation strength, and diffusion fingerprints. This contributes to our understanding of the biased activation of HER2 heterodimers, and the approach is transferable to other membrane receptors targeted by multiple ligands.

## Acknowledgments

We are grateful to Prof. Jacob Piehler for generously providing the plasmid for mEGFP-TMD and for helpful discussions. We thank Petra Freund for assistance in cell culture and Yunqing Li and Sinem Bulmus for helpful discussions.

## Author Contributions

M.H. designed the research. L.T., M.O., and J.K. produced and labeled the nanobody. C.C. performed live-cell single-molecule imaging experiments with support from M.S.D. J.V.R. programmed the analysis software. C.C. performed image analysis with support from J.V.R. and M.S.D. C.C., M.S.D., and M.H. wrote the manuscript with contributions from all authors.

## Data Availability

The datasets generated during the current study are available in the EMBL BioImaging Archive, https://www.ebi.ac.uk/biostudies/bioimages/studies/S-BIAD597.

