## Supplementary Material for "Biased activation of the receptor tyrosine kinase HER2"

**Supplemental Note 1**

Amino acid sequence of HER2-specific nanobody 2Rs15d:

1 QVQLQESGGG SVQAGGSLKL TCAASGYIFN SCGMGWYRQS PGRERELVSR
60 ISGDGDTWHK ESVKGRFTIS QDNVKKTLYL QMNSLKPEDT AVYFCAVCYN
110 LETYWGQGTQ VTVSSAAAXG HHHHHH

with X being the unnatural amino acid 4-azido-L-phenylalanine.= UAA

**Supplemental Note 2**

We compared our finding with receptor expression levels in HeLa cells. RNA expression levels were compared from the Human Protein Atlas v21.1 (https://www.proteinatlas.org/, **Figure S9A**) (Uhlén et al. 2015). RNA-seq data indicated the highest expression level for HER2 in HeLa cells, closely followed by that of EGFR. Comparably, RNA expression levels for HER3 and HER4 are relatively low, although that of HER4 is found at a slightly higher value. The protein expression levels of the ErbB family in HeLa cells report the highest expression level for EGFR, with slightly lower expression levels for HER2 and HER4, while data for HER3 is not available (https://www.proteomicsdb.org/, **Figure S9B**) (Schmidt et al. 2018). Other studies reported no detectable HER3 receptors on the plasma membrane of HeLa cells using fluorescence-activated cell sorting (FACS) (Chen B, Mao R, Wang H, She J. 2010) or confocal microscopy (Belleudi et al. 2012).

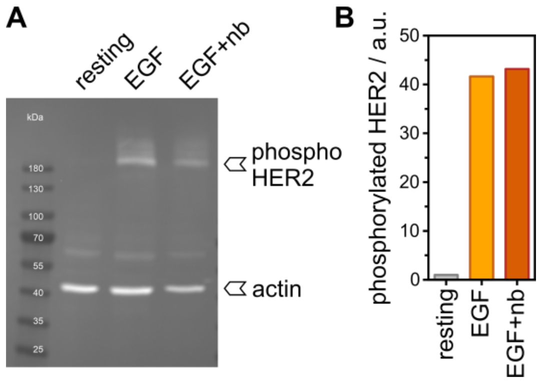


**Supplementary Figure S1. HER2 phosphorylation in the presence and absence of the HER2 nanobody.**
(A) Western blot of cell lysate from HeLa cells in resting state, and treated with either 20 nM EGF or 20 nM EGF and 2 nM anti-HER2 nanobody, stained for phosphorylated HER2. (B) Analysis of the band intensities of phosphorylated HER2 as shown in A. The addition of the nanobody does not impair the ability of HER2 to get cross-phosphorylated after ligand stimulation of the cells. Band intensities were normalized against the housekeeping gene.

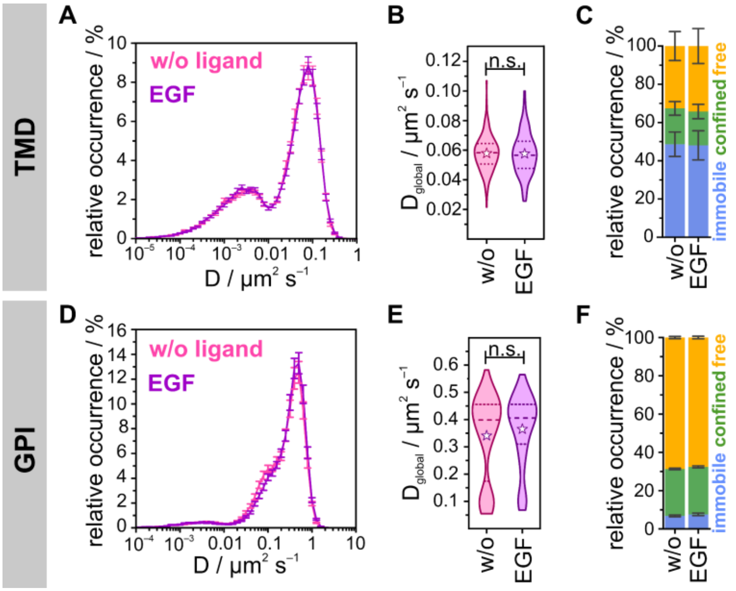


**Supplementary Figure S2. Diffusion dynamics of mEGFP-TMD labeled with AbberiorStar635P-labeled anti-GFP nanobody and GPI-mEos3.2 in untreated and EGF-treated HeLa cells**

(A) Distribution of the diffusion coefficients of the TMD from uPAINT experiments for resting (pink, $D_{global}$ = 0.058 ± 0.004 µm$^2$ s$^{-1}$) and EGF-treated (lilac, $D_{global}$ = 0.058 ± 0.004 µm$^2$ s$^{-1}$) living HeLa cells (N = 140) at 22 °C.

(B) Global diffusion coefficients are displayed as violin plots with dotted lines marking the quartiles, dashed lines the median, and stars representing mean values (p = 0.696).

(C) Relative occurrences of immobile, confined, and freely diffusing particles.

(D) Distribution of the diffusion coefficients of GPI-anchored mEos3.2 from sptPALM experiments for resting (pink, $D_{global}$ = 0.342 ± 0.013 µm$^2$ s$^{-1}$) and EGF-treated (lilac, $D_{global}$ = 0.366 ± 0.017 µm$^2$ s$^{-1}$) living HeLa cells (N = 160) at 23 °C.

(E) Global diffusion coefficients displayed as violin plots with dotted lines marking the quartiles, dashed lines the median, and stars representing mean values (p = 0.360).

(F) Relative occurrences of immobile, confined and freely diffusing particles.

Error bars are defined by SEMs; p > 0.05 no significant difference (n.s.).

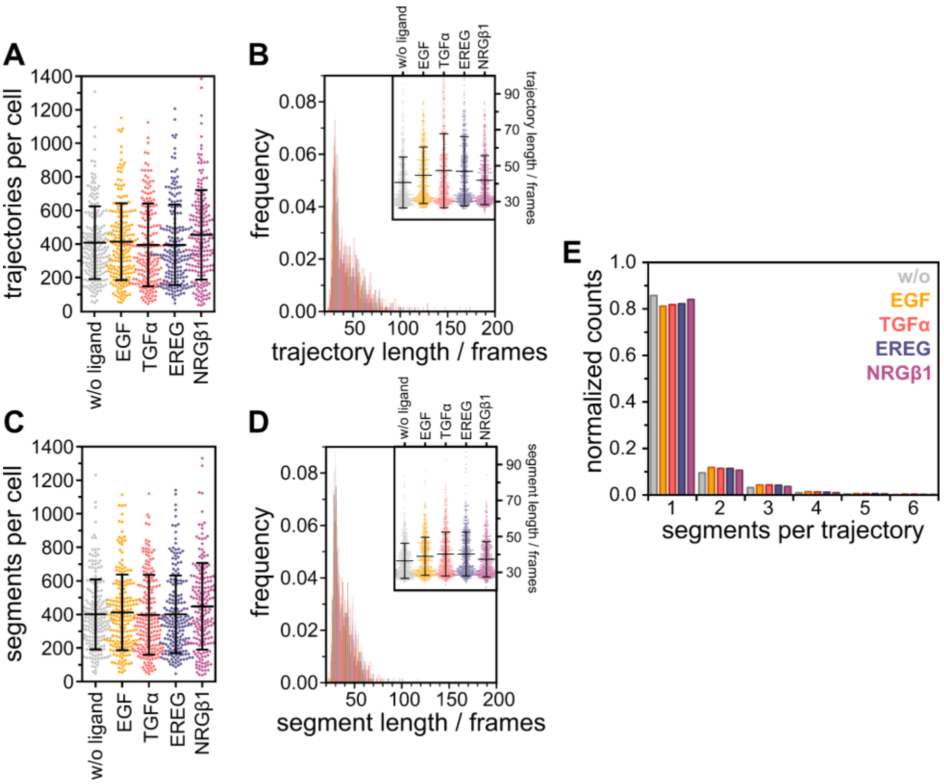


**Supplementary Figure S3. Analysis of segments and trajectories per cell**

(A) Diamonds indicate the mean number of trajectories per cell with the interval representing the overall mean value ± its standard deviation. Overall mean values per condition are 418 ± 222 for resting HER2, 423 ± 232 for EGF, 402 ± 250 for TGFα, 388 ± 234 for EREG, and 466 ± 270 for NRGβ1 stimulated HER2.

(B) Mean trajectory length per cell plotted as a histogram and overlaid for all four conditions. The inlay displays the distribution of trajectory lengths per cell. Data points indicate the mean trajectory length per cell with the interval representing the overall mean value ± its standard deviation. Overall mean values per condition are 40 ± 8 for resting HER2, 44 ± 9 for EGF, 46 ± 12 for TGFα, 44 ± 11 for EREG, and 41 ± 8 for NRGβ1 stimulated cells.

(C) Diamonds indicate the mean number of analyzed segments per cell with the interval representing the overall mean value ± its standard deviation. Overall mean values per condition are 409 ± 212 for resting HER2, 419 ± 227 for EGF, 405 ± 242 for TGFα, 407 ± 234 for EREG, and 457 ± 262 for NRGβ1 stimulated cells.

(D) The mean segment length per cell was plotted as a histogram and overlaid for all four conditions. The inlay displays the distribution of segment lengths per cell. Data points indicate the mean segment length per cell with the interval representing the overall mean value ± its standard deviation. Overall mean values per condition are 36 ± 6 for resting HER2, 39 ± 6 for EGF, 40 ± 7 for TGFα, 40 ± 7 for EREG, and 37 ± 5 for NRGβ1 stimulated cells.

(E) The mean number of segments per trajectory is plotted as a histogram with ~85% of trajectories containing only one segment.

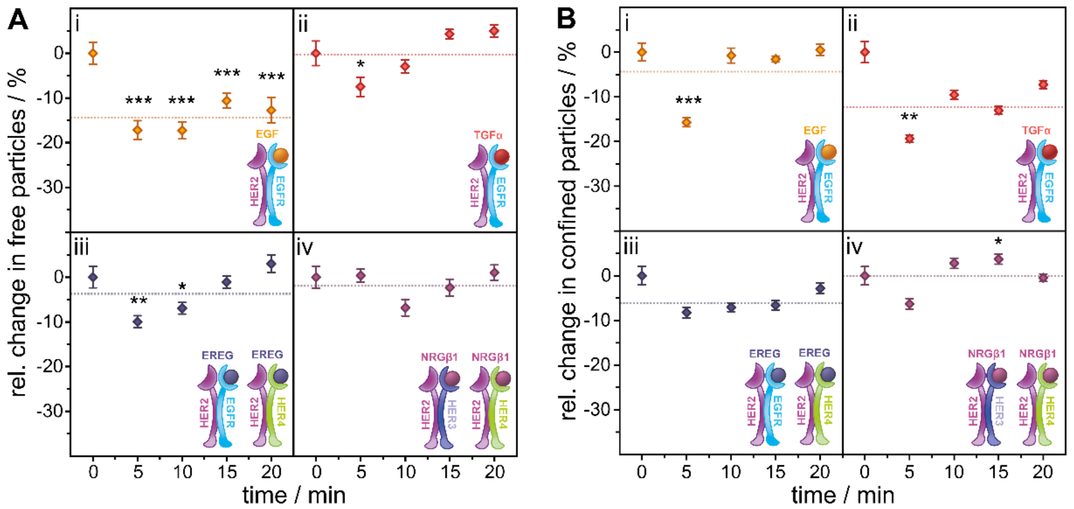


**Supplementary Figure S4. Relative change in the temporal response of freely and confined moving HER2 in living HeLa cells upon ligand stimulation compared to the resting condition**

Related to Figure 3A.

(A) Relative change of molecules classified as freely diffusing plotted against the duration of ligand stimulation.

(B) Relative change of confined receptors plotted against the duration of ligand stimulation.

Relative changes were calculated from mean values of 40 cells per interval. Receptor models indicate the expected ligand-orchestrated interactions between HER2 and other receptors of the family. The dotted lines represent mean values of the relative change over the time of ligand stimulation. Error bars in dot plots represent the standard error of the difference (SED); for bar plots, the standard error of the mean (SEM) is depicted. Significance was tested for stimulated cells vs. resting cells of the same samples before calculating the relative change with $p > 0.05$ no significant difference (no label), $p < 0.05$ significant difference (*), $p < 0.01$ very significant difference (**), $p < 0.001$ highly significant difference (***) between means.

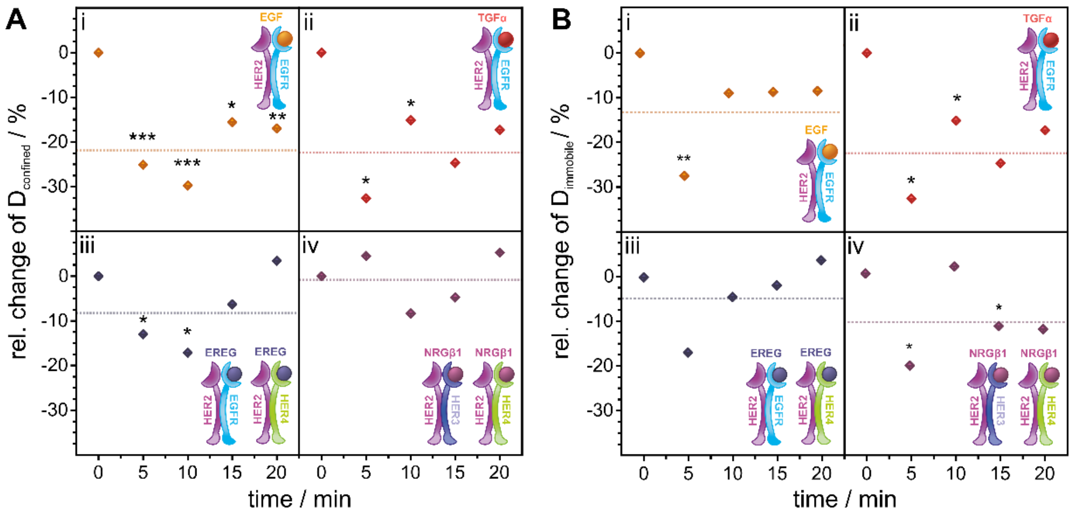


**Supplementary Figure S5. Relative change in the temporal response of diffusion coefficients of confined and immobile HER2 in living HeLa cells upon ligand stimulation compared to the resting condition**

Related to Figure 3B.

(A) Relative change of the diffusion coefficient of confined moving particles plotted against the duration of ligand stimulation.

(B) Relative change of the diffusion coefficient of immobile particles plotted against the duration of ligand stimulation.

Relative changes were calculated from mean values of 40 cells per interval. Receptor models indicate the expected ligand-orchestrated interactions between HER2 and other receptors of the family. The dotted lines represent mean values of the relative change over the time of ligand stimulation. Error bars in dot plots represent the standard error of the difference (SED); for bar plots, the standard error of the mean (SEM) is depicted. Significance was tested for stimulated cells vs. resting cells of the same samples before calculating the relative change with $p > 0.05$ no significant difference (no label), $p < 0.05$ significant difference (*), $p < 0.01$ very significant difference (**), $p < 0.001$ highly significant difference (***) between means.

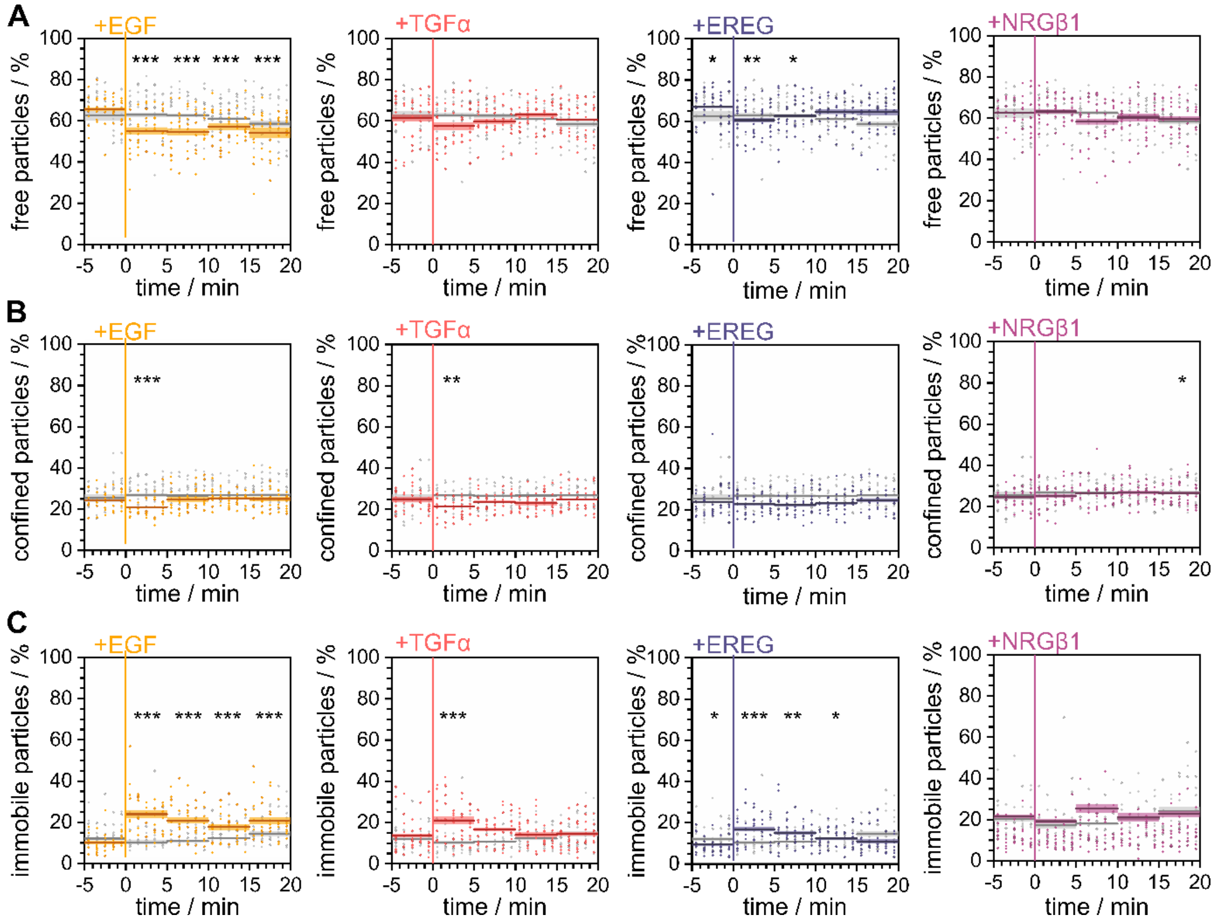


**Supplementary Figure S6. Temporal change of the relative occurrence of diffusion modes**

Related to Figures 3A and S3.

(A) The uncorrected percentages of freely and (B) confined moving as well as (C) immobile HER2 after ligand stimulation in comparison with the resting condition in living HeLa cells (N = 200) are plotted against the time. Data points represent mean values per cell, lines indicate mean values over 5 min with confidence bands representing the SEM. Significance was tested for stimulated cells vs. untreated cells from the same sample. For time windows from –5 to 0 min, the significance was tested between the respective samples and the resting-only condition. Ligand was added at time point 0.

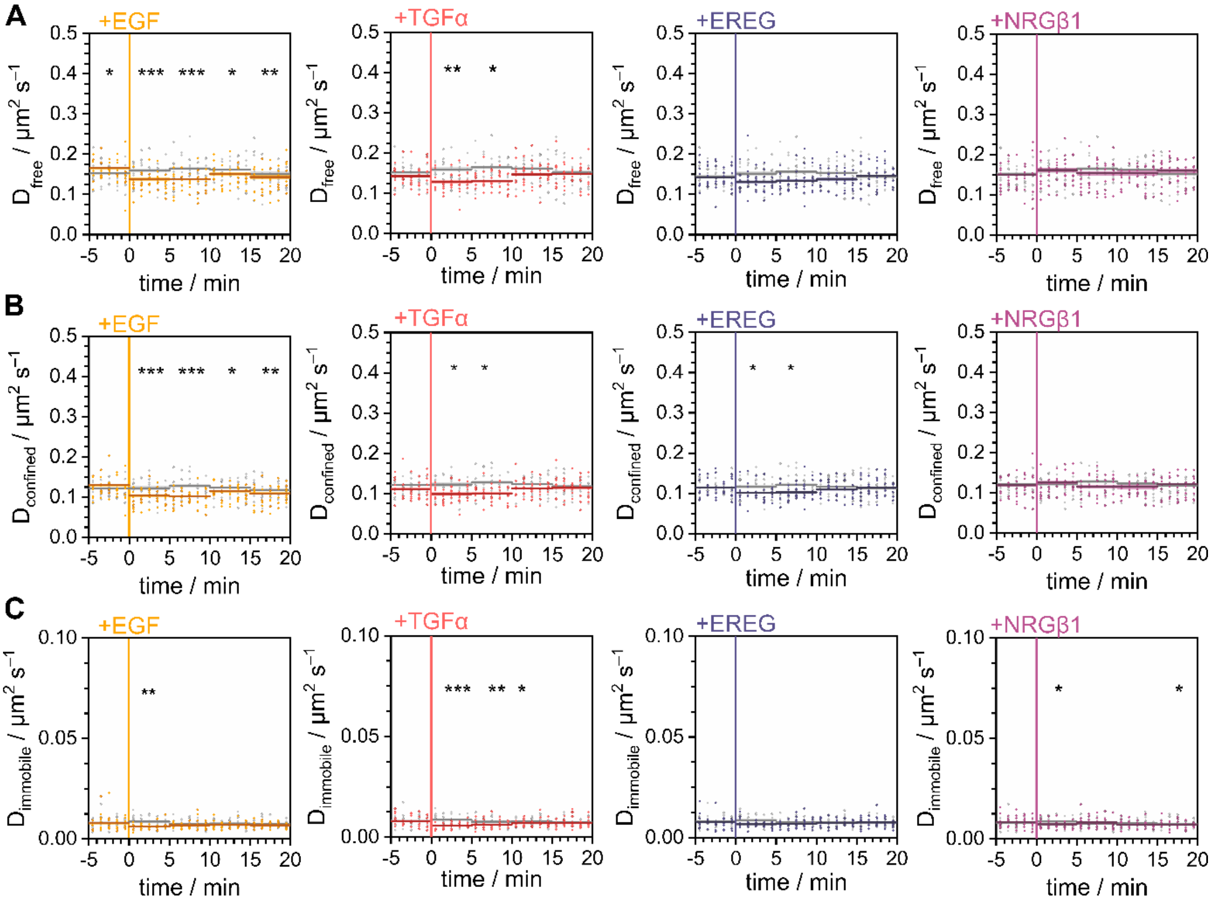


**Supplementary Figure S7. Temporal change of the diffusion coefficients**

Related to Figures 3B and S4.

Diffusion coefficients of (A) freely and (B) confined moving as well as (C) immobile HER2 after ligand stimulation plotted in comparison with the resting condition in living HeLa cells (N = 200). Data points represent mean values per cell, lines indicate mean values over 5 min with confidence bands representing the SEM. Significance was tested for stimulated cells vs. untreated cells from the same sample. For time windows from –5 to 0 min significance was tested between the respective samples and the resting-only condition. Ligand was added at time point 0.

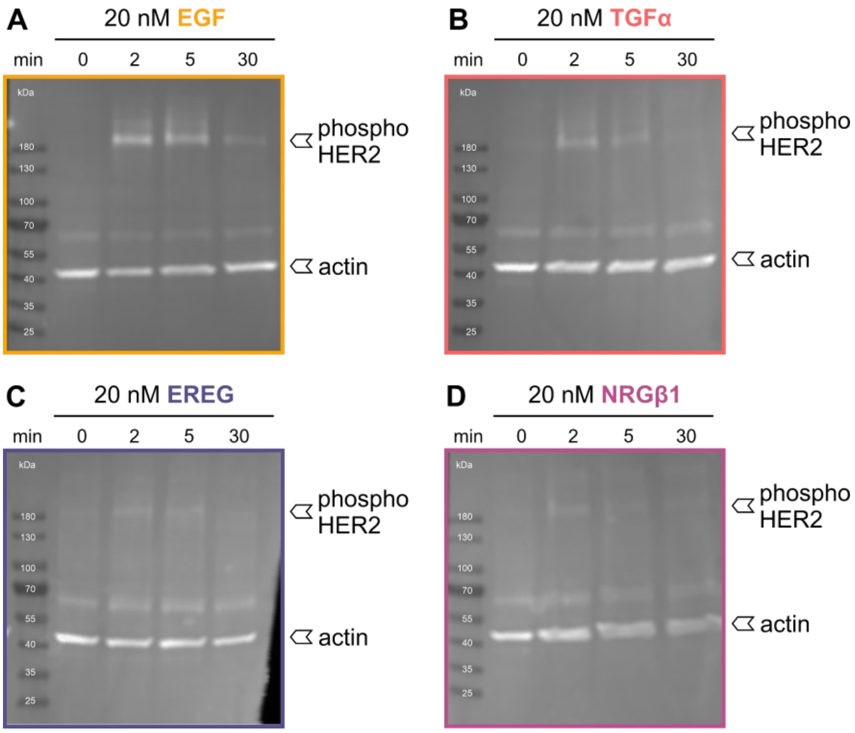


**Supplementary Figure S8. Western blot analysis of ligand-induced phosphorylation of HER2 in HeLa cells**

Related to Figure 3.

An antibody against the phosphorylated tyrosine residues Y1221/1222 of HER2 was applied next to an anti-actin antibody labeling actin as a housekeeping gene. Page ruler served as a size marker on all blots. Tyrosine 1221/1222 was chosen as the target as this site serves as a junction to the Ras-Raf-MAPK pathway.

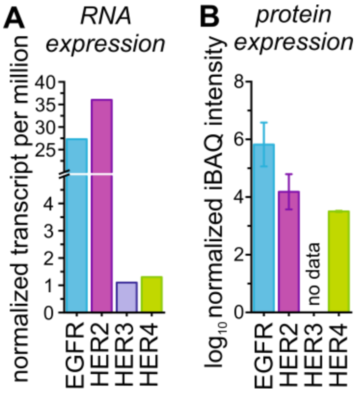


**Supplementary Figure S9. RNA and protein expression levels of ErbB receptors**

(A) RNA expression levels in normalized transcripts per million are taken from https://www.proteinatlas.org/.

(B) Protein expression levels obtained by applying the intensity-based absolute quantification (iBAQ) algorithm were taken from https://www.proteomicsdb.org/.

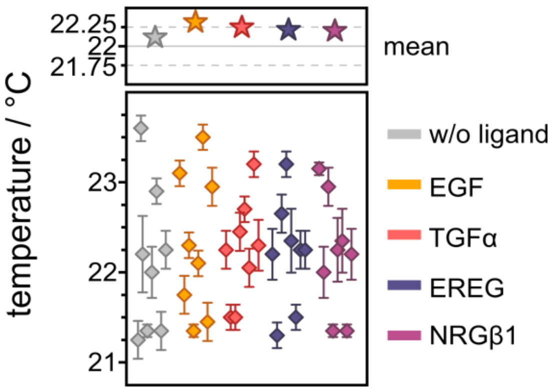


**Supplementary Figure S10. Mean temperature per sample**

Temperatures are color-coded with the respective ligand used for stimulation during that specific measurement. Diamonds represent mean temperatures per coverslip with the respective SEM. Stars represent overall mean values per condition.

**Supplementary Table S1. Diffusion coefficients of mEGFP-TMD in unstimulated and EGF-stimulated cells**

Global diffusion coefficients, and diffusion coefficients for the different diffusion types, are given as mean values derived from 140 cells (errors represent SEMs).

| Sample | $D_{global}$ / µm$^2$s$^{-1}$ | $D_{immobile}$ / µm$^2$s$^{-1}$ | $D_{confined}$ / µm$^2$s$^{-1}$ | $D_{free}$ / µm$^2$s$^{-1}$ |
|---|---|---|---|---|
| **w/o ligand** | 0.058 ± 0.004 | 0.0039 ± 0.0006 | 0.062 ± 0.007 | 0.092 ± 0.005 |
| **EGF** | 0.058 ± 0.004 | 0.0038 ± 0.0005 | 0.064 ± 0.008 | 0.093 ± 0.005 |

**Supplementary Table S2. Mann-Whitney-U test for comparison of global, immobile, confined, and freely diffusing mEGFP-TMD molecules of unstimulated cells with EGF stimulated cells**

*z*-scores derived from the *U*-statistic are listed next to the corresponding *p*-values and assigned levels of significance (*LOS*). Significance level α = 0.05, p ≥ 0.05 no significant difference (n.s.) p < 0.05 significant difference (*), p < 0.01 very significant difference (**), p < 0.001 highly significant difference (***).

| Sample 1 | Sample 2 | global | | | immobile | | | confined | | | free | | |
|---|---|---|---|---|---|---|---|---|---|---|---|---|---|
| | | *z* | *p* | *LOS* | *z* | *p* | *LOS* | *z* | *p* | *LOS* | *z* | *p* | *LOS* |
| **w/o ligand** | **EGF** | 0.39 | 0.70 | n.s. | 1.05 | 0.29 | n.s. | –1.99 | 0.046 | * | –0.34 | 0.74 | n.s. |

**Supplementary Table S3. Percentage of mEGFP-TMD molecules assigned to the three diffusion types immobile, confined, and free in unstimulated and EGF-stimulated cells**
Mean values for 140 cells are listed with their respective SEMs for each diffusion type.

| Sample | immobile / % | confined / % | free / % |
|---|---|---|---|
| **w/o ligand** | 33 ± 7 | 19 ± 4 | 49 ± 6 |
| **EGF** | 34 ± 9 | 18 ± 4 | 48 ± 8 |

**Supplementary Table S4. Mann-Whitney-U test for comparison of the percentage of immobile, confined, and freely diffusing TMD-mEGFP molecules in unstimulated and ligand-stimulated cells**
*z*-scores derived from the *U*-statistic are listed next to the corresponding *p*-values and assigned levels of significance (*LOS*). Significance level $\alpha = 0.05$, $p \geq 0.05$ no significant difference (n.s.) $p < 0.05$ significant difference (*), $p < 0.01$ very significant difference (**), $p < 0.001$ highly significant difference (***).

| Sample 1 | Sample 2 | immobile | | | confined | | | free | | |
|---|---|---|---|---|---|---|---|---|---|---|
| | | *z* | *p* | *LOS* | *z* | *p* | *LOS* | *z* | *p* | *LOS* |
| **w/o ligand** | **EGF** | –1.69 | 0.09 | n.s. | 2.24 | 0.02 | * | 0.78 | 0.43 | n.s. |

**Supplementary Table S5. Diffusion coefficients of GPI-mEos3.2 in unstimulated and EGF-stimulated cells**
Global diffusion coefficients, and diffusion coefficients for the different diffusion types, are given as mean values derived from 160 cells (errors represent SEMs).

| Sample | $D_{global}$ / µm²s⁻¹ | $D_{immobile}$ / µm²s⁻¹ | $D_{confined}$ / µm²s⁻¹ | $D_{free}$ / µm²s⁻¹ |
|---|---|---|---|---|
| **w/o ligand** | 0.342 ± 0.013 | 0.381 ± 0.015 | 0.30 ± 0.02 | 0.006 ± 0.002 |
| **EGF** | 0.367 ± 0.017 | 0.42 ± 0.02 | 0.33 ± 0.03 | 0.006 ± 0.002 |

**Supplementary Table S6. Mann-Whitney-U test for comparison of global, immobile, confined, and freely diffusing GPI-mEos3.2 molecules of unstimulated cells with EGF-stimulated cells**

*z*-scores derived from the *U*-statistic are listed next to the corresponding *p*-values and assigned levels of significance (*LOS*). Significance level α = 0.05, p ≥ 0.05 no significant difference (n.s.) p < 0.05 significant difference (*), p < 0.01 very significant difference (**), p < 0.001 highly significant difference (***).

| Sample 1 | Sample 2 | global | | | immobile | | | confined | | | free | | |
|---|---|---|---|---|---|---|---|---|---|---|---|---|---|
| | | *z* | *p* | *LOS* | *z* | *p* | *LOS* | *z* | *p* | *LOS* | *z* | *p* | *LOS* |
| **w/o ligand** | **EGF** | –0.92 | 0.36 | n.s. | 0.30 | 0.77 | n.s. | –1.33 | 0.18 | * | –1.22 | 022 | n.s. |

**Supplementary Table S7. Percentage of GPI-mEos3.2 molecules assigned to the three diffusion types immobile, confined, and free in unstimulated and EGF-stimulated cells**

Mean values for 160 cells are listed with their respective SEMs for each diffusion type.

| **Sample** | **immobile / %** | **confined / %** | **free / %** |
|---|---|---|---|
| **w/o ligand** | 6.8 ± 0.4 | 24.5 ± 0.4 | 68.7 ± 0.6 |
| **EGF** | 7.6 ± 0.7 | 24.8 ± 0.5 | 67.6 ± 0.7 |

**Supplementary Table S8. Mann-Whitney-U test for comparison of the percentage of immobile, confined, and freely diffusing GPI-mEos3.2 molecules in unstimulated and ligand-stimulated cells**

*z*-scores derived from the *U*-statistic are listed next to the corresponding *p*-values and assigned levels of significance (*LOS*). Significance level α = 0.05, p ≥ 0.05 no significant difference (n.s.) p < 0.05 significant difference (*), p < 0.01 very significant difference (**), p < 0.001 highly significant difference (***).

| Sample 1 | Sample 2 | immobile | | | confined | | | free | | |
|---|---|---|---|---|---|---|---|---|---|---|
| | | *z* | *p* | *LOS* | *z* | *p* | *LOS* | *z* | *p* | *LOS* |
| **w/o ligand** | **EGF** | –0.60 | 0.55 | n.s. | –0.30 | 0.77 | n.s. | 0.88 | 0.37 | n.s. |

**Supplementary Table S9. Binding of EGF family ligands to HER receptors summarized from published data**

Interactions with monomeric receptors and heterodimers are listed. The relative strength of ligand binding/activation to receptors is listed in the right column while the relative strength in activation is listed in the lowest row.

| **Receptor / Ligand** | **EGF** | **TGFα** | **EREG** | **NRGβ1** | **Relative Binding Affinity** |
|---|---|---|---|---|---|
| **EGFR** | x[a,b] | x[a,b] | x[a,b] | | EGF > TGFα > EREG[a] |
| **HER3** | | | | x[a,b] | |
| **HER4** | | | x[b] | x[a,b] | |
| **HER2:EGFR** | x[a] | x[a] | x[a,d] | | EGF > TGFα > EREG[a] |
| **HER2:HER3** | | | x[a,c,d] | x[a] | NRGβ1 > EREG[a] |
| **HER2:HER4** | x[a] | x[a] | x[a,d] | x[a] | NRGβ1 > EGF > EREG, TGFα[a] |
| **Relative Activation Strength** | 1:2 > 1:4[a] | 1:2 >> 2:4[d] | 1:2 > 2:3, 2:4[d] | HER4 > HER3[e,f] 1:3 > 1:4[a] | |

[a] Jones 1999 (10.1016/s0014-5793(99)00283-5)
[b] Wilson 2008 (10.1016/j.pharmthera.2008.11.008)
[c] Barber 2019 (10.1093/jnci/djz231)
[d] Shelly 1998 (10.1074/jbc.273.17.10496)
[e] Pinkas-Kramarski 1998 (10.1128/MCB.18.10.6090)
[f] Tzahar 1994 (10.1016/S0021-9258(17)31521-1)

**Supplementary Table S10. Diffusion coefficients of HER2 in unstimulated and ligand-stimulated cells**

Related to Figures 1C,F, 2C,F.

Mean diffusion coefficients for 128 cells are listed next to mean values according to diffusion types with their respective SEMs.

| Sample | $D_{global}$ / µm²s⁻¹ | $D_{immobile}$ / µm²s⁻¹ | $D_{confined}$ / µm²s⁻¹ | $D_{free}$ / µm²s⁻¹ |
|---|---|---|---|---|
| **w/o ligand** | 0.125 ± 0.005 | 0.0079 ± 0.0016 | 0.116 ± 0.009 | 0.150 ± 0.006 |
| **EGF** | 0.102 ± 0.005 | 0.0067 ± 0.0011 | 0.102 ± 0.008 | 0.135 ± 0.006 |
| **TGFα** | 0.111 ± 0.005 | 0.0065 ± 0.0012 | 0.107 ± 0.008 | 0.138 ± 0.005 |
| **EREG** | 0.113 ± 0.005 | 0.0070 ± 0.0014 | 0.107 ± 0.008 | 0.137 ± 0.06 |
| **NRGβ1** | 0.122 ± 0.005 | 0.0074 ± 0.0014 | 0.114 ± 0.008 | 0.149 ± 0.006 |

**Supplementary Table S11. Mann-Whitney-U test for comparison of global, immobile, confined, and free diffusion coefficients of HER2 in unstimulated cells with ligand-stimulated cells**

Related to Figures 1C,F, 2C,F.

*z*-scores derived from the *U*-statistic are listed next to the corresponding *p*-values and assigned levels of significance (*LOS*). Significance level α = 0.05, p ≥ 0.05 no significant difference (n.s.) p < 0.05 significant difference (*), p < 0.01 very significant difference (**), p < 0.001 highly significant difference (***).

| Sample 1 | Sample 2 | global | | | immobile | | | confined | | | free | | |
|---|---|---|---|---|---|---|---|---|---|---|---|---|---|
| | | *z* | *p* | *LOS* | *z* | *p* | *LOS* | *z* | *p* | *LOS* | *z* | *p* | *LOS* |
| **w/o ligand** | **EGF** | 7.3 | $7 \cdot 10^{-13}$ | *** | 5.9 | $4 \cdot 10^{-9}$ | *** | 5.7 | $2 \cdot 10^{-8}$ | *** | 4.7 | $2 \cdot 10^{-6}$ | *** |
| | **TGFα** | 4.5 | $8 \cdot 10^{-6}$ | *** | 6.4 | $1 \cdot 10^{-10}$ | *** | 3.9 | $8 \cdot 10^{-5}$ | *** | 3.7 | $2 \cdot 10^{-4}$ | *** |
| | **EREG** | 4.0 | $7 \cdot 10^{-5}$ | *** | 4.0 | $7 \cdot 10^{-5}$ | *** | 4.0 | $5 \cdot 10^{-5}$ | *** | 4.4 | $1 \cdot 10^{-5}$ | *** |
| | **NRGβ1** | 0.7 | 0.5 | n.s. | 2.6 | $8 \cdot 10^{-3}$ | ** | 1.1 | 0.3 | n.s. | 0.2 | 0.9 | n.s. |

**Supplementary Table S12. Corrected percentage of HER2 molecules assigned to the three diffusion types immobile, confined, and free in unstimulated and ligand-stimulated cells**

Related to Figures 1E, 2E.

Mean values for 128 cells are listed with their respective SEMs for each diffusion type.

| Sample | immobile / % | confined / % | free / % |
|---|---|---|---|
| **w/o ligand** | 12.0 ± 0.4 | 26.5 ± 0.4 | 61.5 ± 0.6 |
| **EGF** | 22.7 ± 0.7 | 25.1 ± 0.4 | 52.2 ± 0.8 |
| **TGFα** | 14.9 ± 0.6 | 23.8 ± 0.4 | 61.3 ± 0.7 |
| **EREG** | 16.5 ± 0.5 | 24.9 ± 0.4 | 58.6 ± 0.6 |
| **NRGβ1** | 12.5 ± 0.5 | 27.1 ± 0.4 | 60.3 ± 0.8 |

**Supplementary Table S13. Mann-Whitney-U test for comparison of the corrected percentage of immobile, confined, and freely diffusing HER2 in unstimulated cells with ligand-stimulated cells**

Related to Figures 1E, 2E.

*z*-scores derived from the *U*-statistic are listed next to the corresponding *p*-values and assigned levels of significance (*LOS*). Significance level α = 0.05, p ≥ 0.05 no significant difference (n.s.) p < 0.05 significant difference (*), p < 0.01 very significant difference (**), p < 0.001 highly significant difference (***).

| Sample 1 | Sample 2 | immobile | | | confined | | | free | | |
|---|---|---|---|---|---|---|---|---|---|---|
| | | *z* | *p* | *LOS* | *z* | *p* | *LOS* | *z* | *p* | *LOS* |
| **w/o ligand** | **EGF** | –11.6 | $3 \cdot 10^{-31}$ | *** | 2.9 | $4 \cdot 10^{-3}$ | ** | 8.7 | $2 \cdot 10^{-19}$ | *** |
| | **TGFα** | –3.3 | $8 \cdot 10^{-4}$ | *** | 4.7 | $3 \cdot 10^{-6}$ | *** | 0.4 | 0.7 | n.s. |
| | **EREG** | –6.9 | $5 \cdot 10^{-12}$ | *** | 3.1 | $2 \cdot 10^{-3}$ | ** | 3.4 | $6 \cdot 10^{-4}$ | *** |
| | **NRGβ1** | –0.7 | 0.49 | n.s. | –1.0 | 0.3 | n.s. | 1.1 | 0.3 | n.s. |
